# Dysregulation of The Chromatin Environment Leads to Differential Alternative Splicing as A Mechanism Of Disease In a Human Model of Autism Spectrum Disorders

**DOI:** 10.1101/2022.09.08.507162

**Authors:** Calvin S. Leung, Shoshanna Rosenzweig, Brian Yoon, Nicholas A. Marinelli, Ethan W. Hollingsworth, Abbie M. Maguire, Mara M. Cowen, Michael Schmidt, Jaime Imitola, Ece D. Gamsiz Uzun, Sofia B. Lizarraga

## Abstract

Autism spectrum disorder (ASD) affects 1 in 44 children. Chromatin regulatory proteins are overrepresented among genes that contain high risk variants in ASD. Disruption of the chromatin environment leads to widespread dysregulation of gene expression, which is traditionally thought as a mechanism of disease pathogenesis associated with ASD. Alternatively, alterations in chromatin dynamics could also lead to dysregulation of alternative splicing, which is understudied as a mechanism of ASD pathogenesis. The anticonvulsant valproic acid (VPA) is a well-known environmental risk factor for ASD that acts as a class I histone deacetylase (HDAC) inhibitor. However, the precise molecular mechanisms underlying defects in human neuronal development associated with exposure to VPA are understudied. To dissect how VPA exposure and subsequent chromatin hyperacetylation influence molecular signatures involved in ASD pathogenesis, we conducted RNA sequencing (RNA-seq) in human cortical neurons that were treated with VPA. We observed that differentially expressed genes (DEGs) were enriched for mRNA splicing, mRNA processing, histone modification, and metabolism related gene sets. Furthermore, we observed widespread increase in the number and the type of alternative splicing events. Analysis of differential transcript usage (DTU) showed that exposure to VPA induces extensive alterations in transcript isoform usage across neurodevelopmentally important genes. Finally, we find that DEGs and genes that display DTU overlap with known ASD-risk genes. Together, these findings suggest that, in addition to differential gene expression, changes in alternative splicing correlated with alterations in the chromatin environment could act as an additional mechanism of disease in ASD.

## INTRODUCTION

Autism spectrum disorders (ASD) are highly heritable neurodevelopmental disorders with complex genetic etiology ^1^. Several lines of evidence suggests that gene-environment interaction could influence the risk of developing ASD and contribute to its complex etiology ^2,3^. Valproic acid (VPA) is an anticonvulsant drug used in the treatment of epilepsy and is a well-characterized environmental risk factor for ASD ^4^. An increasing number of studies have shown that prenatal exposure to VPA during pregnancy increased the risk of ASD ^4–8^. Exposure to VPA during pregnancy has also been associated with the fetal valproate syndrome (FVS), characterized by a higher incidence of neural tube defects, craniofacial abnormalities, developmental delay, intellectual disability, and ASD ^9,10^. However, mechanistic studies that interrogate the association between VPA exposure and increased ASD risk, or neurodevelopmental defects in humans, are lacking.

Acetylation of histones by histone acetyltransferases (HATs) results in a dispersed or ‘open’ chromatin conformation, allowing physical access for transcription factors to bind to DNA and generally activates transcription ^11,12^. Conversely, histone deacetylases (HDACs) remove acetyl groups from histones, compacting chromatin and in general are thought to lead to transcriptional repression^12^. VPA was identified as a class I selective HDAC inhibitor and has also been found to induce proteasomal degradation of HDAC2 ^13,14^. VPA’s HDAC inhibition has been shown to be relevant to ASD pathogenesis, because a VPA analog that lacks the HDAC inhibitory function does not cause ASD-like behaviors in mice ^15^. HDAC inhibition leads to hyperacetylation of histones H3 and H4 and results in transcriptional dysregulation^16,17^. In addition to its role in regulating transcription, histone acetylation has also been shown to contribute to the regulation of RNA splicing, an essential process of eukaryotic gene expression ^18–20^. Increasing evidence suggest that a dysregulated chromatin environment caused by HDAC inhibition modulates alternative splicing ^21,22^. Therefore, HDAC inhibition by VPA could alter both gene expression and alternative splicing, which suggests a plausible molecular mechanism underlying VPA’s role as an environmental risk factor for ASD. Taken together, these findings suggest the potential involvement of chromatin regulatory mechanisms in the underlying pathology of ASD associated with VPA exposure.

Epidemiological and animal studies suggest a strong association between prenatal exposure to VPA during the first gestational trimester higher incidence of ASD and birth defects ^6,23,24^. At this developmental stage, the cerebral cortex is the most rapidly developing structure in the brain. Development of the cerebral cortex requires the careful coordination of progressive complex cellular events such as neuronal progenitor proliferation, neuronal migration, generation of neurons, neuronal morphogenesis and synapse development ^25^. Even slight alterations in the timing of these processes could impact cortical development, and lead to deficits in cognition and behavior ^26^. The advent of human stem cell derived neural models constitutes an unprecedented opportunity to study the molecular mechanisms underlying the early stages of human brain development that would be otherwise inaccessible ^27–29^. To investigate the effect of VPA exposure on human neuronal development and identify molecular mechanisms underlying ASD pathogenesis, we used male human induced pluripotent stem cells (iPSCs) of a neuro-typical individual with no known history of neuropsychiatric disorders to derive forebrain cortical excitatory neurons ^30^. We used a dual-SMAD inhibition protocol to generate cortical excitatory neurons as it recapitulates the temporal sequence in which neurons arise during development ^30^. Therefore, this approach represents a robust system to analyze transcriptomic changes during early human fetal development ^31^.

To determine the transcriptomic changes that occur with VPA exposure, we performed RNA-sequencing (RNA-seq) in iPSC-derived neurons treated with VPA for 24 hours at day 65 of neuronal induction. Our analysis revealed 6,208 differentially expressed genes (DEGs). Gene set enrichment analysis (GSEA) revealed that these DEGs were enriched for mRNA splicing, mRNA processing, histone modification, and metabolism-related gene sets. Remarkably, we identified a distinct signature of differential spliced genes in neurons treated with VPA that implicates dysregulation of RNA splicing as a mechanism of ASD pathogenesis. Subsequent differential transcript usage (DTU) analysis identified 3,848 genes that showed significant DTU events. We find that genes with DTU shared little overlap with DEGs and were enriched for neurodevelopmental disorder-related genes. However, we found that both DEGs and genes that display DTU largely overlap with known ASD risk genes. Considering the large numbers of chromatin regulators in which high and strong risk variants have been associated with ASD which include histone acetylation modifiers, dysregulation of alternative splicing could be a common mechanism of disease in ASD ^32^. In fact, transcriptomic studies of different subsets of ASD postmortem brain tissue point to an overrepresentation of differential splicing events in patient samples compared to controls ^33,34^. However, the extent to which changes in alternative splicing occur at early stages of human neuronal development in relation to specific ASD genetic or environmental risk factors is understudied. Here we show for the first time that exposure to a well-known environmental risk factor of ASD at early stages of human cortical development, leads to changes in gene expression and alternative splicing that may act synergistically as a mechanism underpinning ASD pathogenesis.

## MATERIALS AND METHODS

### Induced pluripotent stem cell culture

We used a control iPSC line I-93-7 obtained from a healthy neurotypical male individual that was previously characterized ^35^. Human iPSCs were cultured as previously described ^35,36^. Briefly, cells were cultured using a feeder-free system and mTeSR media (STEMCELL technologies catalog # 85850). Cultures were inspected daily for spontaneous differentiating colonies which were manually removed to ensure the purity of the stem cell cultures. iPSCs were passaged every 4 to 7 days with ReLeSR reagent (STEMCELL technologies catalog #100-0484) at a ratio of 1:6 per culture. In order to ensure that no mutations were introduced in the iPSC cultures, we used cultures that were passaged for less than 40 cycles.

### Neuronal differentiation and VPA treatment

Human control iPSCs were differentiated using a previously described dual SMAD inhibition protocol that promotes the formation of the neuroectoderm by preventing the formation of the mesoderm and endoderm by targeting the BMP signaling pathway ^30^. This protocol primarily produces excitatory forebrain cortical neurons in a monolayer and was conducted as previously described ^35^. In brief, human iPSCs were dissociated into single cells using Gentle Cell Dissociation Reagent (STEMCELL Technologies catalog # 100-0485) and were grown for 1 day with STEMdiff Neuronal induction medium (STEMCELL Technologies catalog # 05835), supplemented with 10 μM ROCK inhibitor (Y-27632 dihydrochloride, TOCRIS catalog # 1254). After a confluent monolayer formed, cells were switched to a neuronal induction medium containing two SMAD inhibitors: 1μM Dorsomorphin (STEMGENT catalog # 04-0024) and 10μM SB 431542 (TOCRIS catalog # 1614), and all subsequent steps were conducted as previously described ^30,35^. After day 45 of neuronal induction, cultures were supplemented with 10ng/μl laminin every other day to ensure they remained attached. At day 64, three independent cultures were treated with 200μg/ml VPA or DMSO at day 64 of neuronal induction and were harvested after 24hr for RNA extraction.

### Library preparation and sequencing

To analyze changes in the neuronal transcriptome, cells were harvested on ice, and RNA was extracted on the same day using mirVANA RNA isolation kit (Thermofisher catalog # 1560) per manufacturer’s guidelines. RNA quality was assessed using a Bioanalyzer (Supplementary Figure S1A-B). Samples with RNA integrity number (RIN) higher than 9 were used for RNA sequencing. cDNA libraries were prepared using the Illumina Tru-Seq kit (Illumina catalog #FC-122-1001). Isolated cDNA libraries were sequenced by paired-end chemistry via Illumina HiSeq 2500. On average, 300 millions of 50 bp paired-end reads were obtained from each library.

### RNA-seq data processing and analysis

The quality of raw sequence data in FastQ files generated following sequencing was accessed using FastQC (version 0.11.9) ^37^ (Supplementary Figure S1C). Raw reads were mapped to the Ensembl reference genome 104 (GRCh38) using the transcript-level quantifier Salmon (version 1.5.2) in mapping-based mode ^38^. An average of 149,416,071.5 fragments were observed with an average mapping rate of 82.96% (Supplementary Table S1). The tximport software package (version 1.20.0) was used to produce count matrices from the transcript-level quantification files produced by Salmon ^39^. Following transcript-to-gene mapping, differentially expressed genes (DEGs) were detected using the DESeq2 software package (version 1.32.0) ^40^. Statistically significant DEGs were identified as DEGs with an adjusted *P* < 0.05 and a fold change ≥ |1.5| (Supplementary Table S2).

### Differential alternative mRNA splicing and transcript usage analysis

Differential transcript usage (DTU) events were identified using DRIMseq software package (version 1.20.0) ^41^. Lowly expressed transcripts were filtered from analysis. Transcripts needed to have at least 10 estimated counts in at least 3 samples. Additionally, transcripts needed to have a minimum proportion of 0.1 in at least 3 samples. Statistically significant DTU events were identified following stageR (version 1.14.0) post-processing analysis as events that has an adjusted *P* < 0.05. Differential alternative splicing events were identified using rMATS (version 4.1.1) ^42^. Statistically significant differential alternative splicing events were identified as events with FDR < 0.05 and IncLevelDifference ≥ |0.1|. Exons with total read counts ≤ 20 were excluded from the analysis. ClusterProfiler (version 4.0.5) was used for functional enrichment analysis (GSEA, GO, and DisGeNET) of DEGs and DTU events. Significant enrichment results were obtained using a *q*-value cutoff of < 0.05.

### Validation of RNA-seq by quantitative PCR (qPCR)

Differential gene expression for a subset of genes was validated using qPCR as previously described ^36^. Primers were purchased in IDT (Supplementary Table S3).

## RESULTS

### VPA elicits a distinct molecular signature in human forebrain cortical excitatory neurons

Weighted gene co-expression analysis studies have identified forebrain cortical neurons as hubs for the expression of genes that have high-risk variants associated with ASD ^43^. Because ASD has a higher incidence in males than in females, we examined the effect of VPA on male human iPSC-derived forebrain cortical neurons that were differentiated for 65 days ^35^. iPSCs were differentiated to cortical forebrain excitatory neurons by inhibiting the BMP and TGF-β signaling pathways to induce the forebrain neuroectodermal lineage (Figure 1A) ^30^. To uncouple the effect of VPA on neuronal progenitors, we used neurons grown for 65 days as they already start displaying synaptic activity at this stage ^28^. We isolated high quality RNA from control and VPA-treated samples that were used to prepare RNA-seq libraries (Supplementary Figure S1). RNA-seq was used to determine the genome-wide effects of VPA exposure. We used the Salmon software package ^38^ to quantify transcript abundance from the RNA-seq reads, allowing for both downstream differential gene expression and differential transcript usage analysis (Figure 1B and Supplementary Table S1). Using the DESeq2 software package, we observed, via principal component analysis (PCA), that the sequenced samples showed distinct clustering of biological replicates by treatment group, suggesting that VPA treatment contributed to a distinct transcriptomic profile compared to untreated neurons (Figure 1C). We identified statistically significant differentially expressed genes (DEGs) with adjusted *P* < 0.05 and FC ≥ |1.5|. We observed widespread and distinct changes in gene expression in the VPA-treated samples compared to control samples with 3545 upregulated and 2663 downregulated DEGs (Figure 1D-F, and Supplementary Table S2). To confirm some of the RNA-seq results we conducted quantitative PCR (qPCR) validation on representative upregulated and downregulated DEGs (Supplementary Figure S2 and Supplementary Table S3).

**Figure 1:**
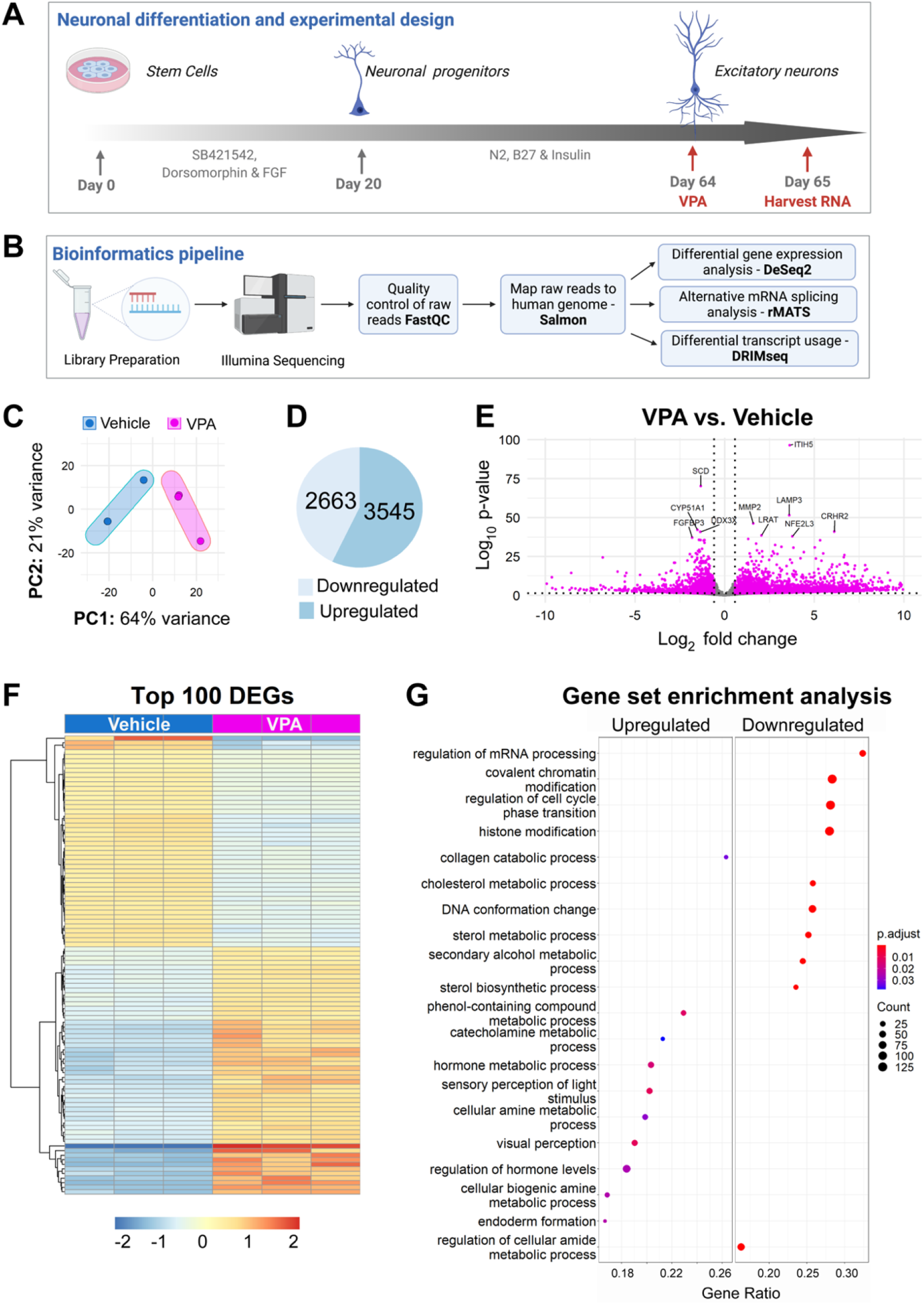
VPA elicits differential gene expression in human cortical excitatory neurons. **(A)** Generation of human cortical excitatory neurons from iPSCs and experimental paradigm. Diagram shows key steps of a double SMAD inhibition protocol used to generate human forebrain cortical excitatory neurons. Addition of dorsomorphin and SB421542 promotes neural fate by blocking the formation of mesoderm and endoderm. Neurons were allowed to mature for 65 days after the initiation of neuronal induction. On Day 64 VPA (1.2 mM) was added to the neurons for 24 h. RNA was extracted for RNA-seq analysis on Day 65. (**B)** Flow chart of the bioinformatics analysis pipeline. RNA libraries were constructed and processed using an Illumina platform. Quality control of raw reads was performed using FastQC. Trimmed reads were mapped to the human genome (GRCh38) using Salmon. Differential gene expression analysis was performed using DESeq2, alternative mRNA splicing analysis was performed using rMATS, and differential transcript usage analysis was performed using DRIMSeq. **(C)** Principal component analysis (PCA) of RNA-seq samples of neurons treated with vehicle (blue) or VPA (magenta). **(D)** Pie chart showing significantly upregulated and downregulated DEGs with cutoffs LFC > |1.5| and adjusted *P* < 0.05. **(E)** Volcano plot showing indicated changes in log_2_ fold changes (LFC) gene expression (x-axis) and −log_10_ *P* values (y-axis) generated from DESeq2 analysis. Vertical dotted lines represent 0.58 LFC (1.5 FC) and the horizontal dotted line represents adjusted *P* = 0.05. Top 10 significant genes based on *P* value are labeled. **(F)** Heat map representation of differential gene expression. The top 100 DEGs were selected based on their adjusted *P* values. Each column represents an independent sample that was either treated with vehicle or treated with VPA. Gene count values were plotted, and color coded to represent expression with red indicating high expression levels and blue indicating low expression levels of individual genes. **(G)** Dot plot displaying GSEA results of upregulated and downregulated DEGs (*P* value cutoff = 0.05). Top 20 enrichment results are shown.

To begin to define the extent to which there were specific molecular signatures altered in response to VPA, we conducted gene set enrichment analysis (GSEA) of DEGs (Supplementary Table S4). Among the topmost enriched gene categories, we found downregulated DEGs that were implicated in the regulation of mRNA processing, covalent chromatin modification, regulation of cell cycle phase transition, and histone modification (Figure 1G). Interestingly, multiple genes encoding mRNA splicing factors were observed within the regulation of mRNA processing pathway. mRNA splicing factors are involved in the formation of mature mRNA through intron excision and exon ligation of precursor mRNA (pre-mRNA). Additionally, alternative mRNA splicing allows for a single gene to generate an average of three alternatively spliced mRNA isoforms, leading to increased complexity of gene expression and protein function in the cell ^44^. Individual gene count analysis of representative alternative splicing factors *PTBP1* (log_2_ FC = −0.6018725), *SRSF12* (log_2_ FC = −1.186584437), and *RBFOX2* (log_2_ FC = −0.5403933) showed consistent downregulation of these genes (Supplementary Figure S3). Downregulation of alternative splicing factors and other mRNA processing genes suggests that VPA may also lead to changes in transcript isoform usage.

In addition to the distinct mRNA processing signature, we identify a chromatin regulatory signature in response to VPA exposure, as genes involved in covalent chromatin modification, histone modification, and DNA conformation change pathways were also found to be downregulated. Genes in these pathways include histone methyltransferases, histone demethylases, histone acetyltransferases, histone deacetylases, and chromatin remodelers. Representative genes in these categories include: *NSD1* (log_2_ FC = −1.225422657), *KDM5B* (log_2_ FC = −1.038048715), *KAT2B* (log_2_ FC = −1.312303), *HDAC6* (log_2_ FC = −0.631808), and *CHD2* (log_2_ FC = −0.9104716) (Supplementary Figure S3). Regulation of the chromatin landscape by chromatin modifiers and/or remodelers has been shown to have widespread effects in the control of gene expression. Furthermore, specific chromatin modifications have also been associated with the control of RNA splicing. Thus, downregulation of chromatin modifiers and remodelers in response to VPA treatment, could have an additive effect in the control of gene expression and alternative splicing, and exacerbate VPA-related phenotypes.

### Profiling of the alternative mRNA splicing landscape in VPA-treated neurons

Dysregulation of histone acetylation by HDAC inhibition has previously been shown to affect alternative RNA splicing ^18,21^. We found downregulation of RNA splicing factors in VPA-treated neurons. Therefore, we sought to determine whether global changes in alternative splicing events were associated with VPA exposure. We conducted replicate multivariate analysis of transcript splicing (rMATS) ^42^, and detected extensive changes in alternative splicing events in the VPA-treated samples after filtering out events with low exonic counts (< 20) and events that showed less than a 10% change (Figure 2A and Supplementary Table S5). Skipped exon (SE) and retained intron (RI) events were the most frequent splicing events detected in VPA-treated neurons at 40% and 22% of total events, respectively (Figure 2B). Other splice events observed were alternative 5’ or 3′ splice site (A5SS or A3SS) and mutually exclusive exon (MXE) (Figure 2B). Interestingly, VPA treatment led to decreased alternative exon inclusion in four out of the five different splice event types, suggesting that histone hyperacetylation could be implicated in the regulation of alternative splicing (Figure 2B). For purpose of illustration of some of the changes we found, we show analysis of two genes *CLASP1* and *CEP170* which were among the most significantly altered in terms of differential splicing. Sashimi plots of representative examples for SE and RI, *CLASP1* and *CEP170*, respectively, showed fewer reads in alternatively spliced exons and greater skipped junction reads in VPA-treated samples (Figure 2C – 2D).

**Figure 2:**
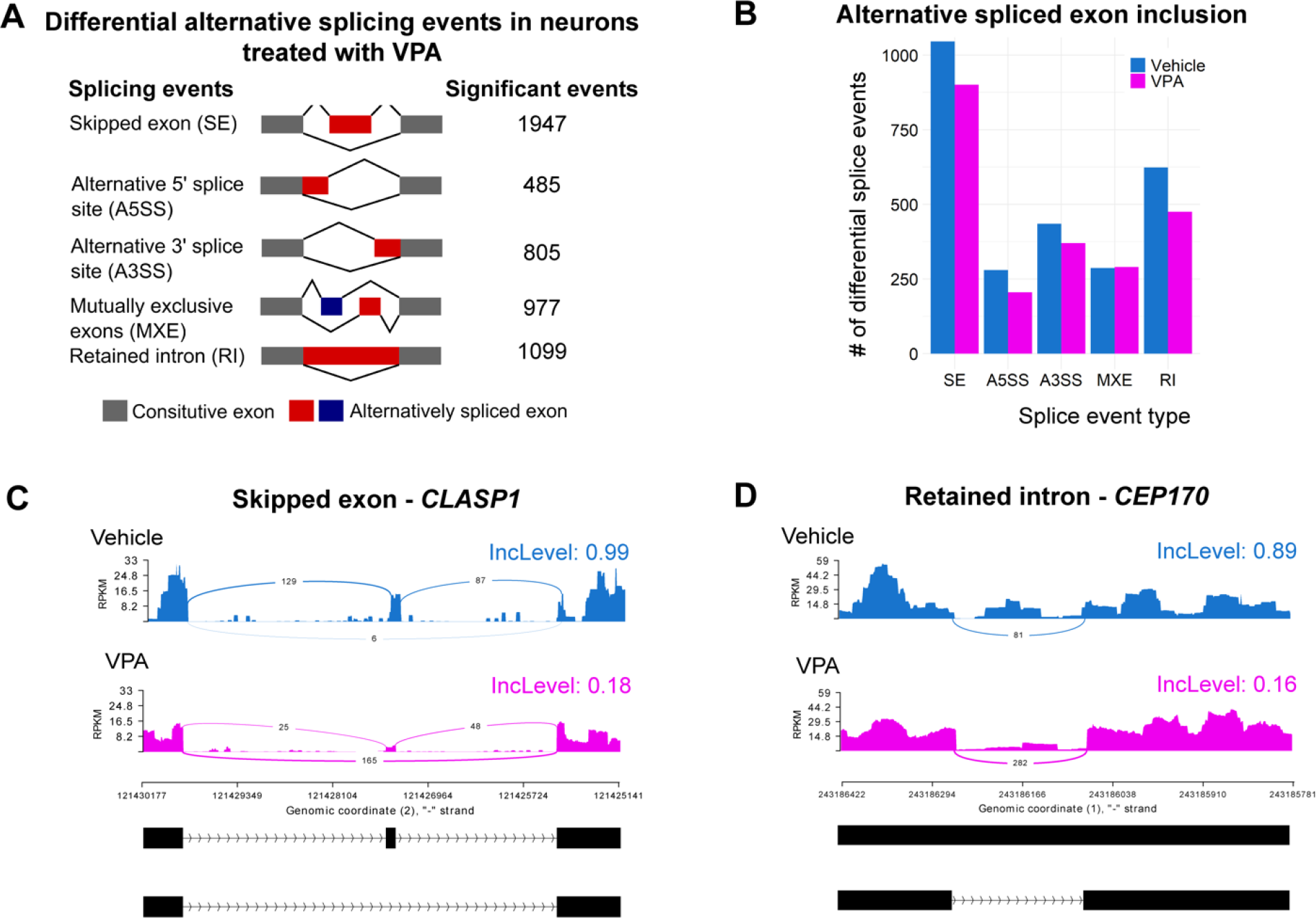
VPA induces widespread changes in alternative mRNA splicing in human neurons. **(A)** Diagram showing the different types of alternative mRNA splicing events identified using rMATS and number of significant events (ΔPercent spliced in (PSI) > 0.1; FDR < 0.05) for each event after filtering (>20 alternative exon counts). **(B)** Bar plot displaying number of significant alternatively spliced exon inclusion events for the different types of alternative mRNA splicing events in neurons treated with vehicle (blue) or VPA (magenta). **(C)** Sashimi plot of the gene *CLASP1* showing a skipped exon event. Alternative exon inclusion levels (IncLevel) for vehicle and VPA are indicated. **(D)** Sashimi plot of *CEP170* shows a retained intron event. Alternative exon inclusion levels (IncLevel) for vehicle and VPA are shown.

### Alterations in transcript usage are observed primarily among genes associated with RNA splicing and genes associated with neurodevelopmental disorders

Differential RNA alternative splicing could have diverse phenotypic effects in the cell due to changes in frequency of non-coding RNAs, mRNA stability, as well as gain or loss of protein domains, all which have the potential to alter cellular function and lead to disease ^45^. Through DRIMSeq differential transcript usage (DTU) analysis ^41^, we analyzed a total of 11,807 genes with a total of 33,633 transcript isoforms (Figure 3A and Supplementary Table S6). We detected extensive changes in DTU events in VPA-treated neurons. Therefore, we sought to define if these changes in transcript isoform expression in VPA-treated neurons compared to control neurons was occurring primarily in the DEGs. We determined the overlap between DEGs with genes that display significant DTU events (Figure 3B). While we found an overlap of 995 DEGs that have DTU, our analysis showed that the majority of DTU (2853) was not occurring in DEGs. To begin to interrogate the extent to which the DTU genes might be different from the DEGs we first conducted gene ontology (GO) biological process analysis on the DTU specific genes (Supplementary Table S7). Interestingly, we identified an overrepresentation of genes involved in RNA biology which included RNA splicing, regulation of RNA stability, and regulation of RNA metabolism (Figure 3C). Next, we characterized the correlation of DTU with specific disease pathways using DISGeneNet databases ^46^ (Supplementary Table S8). This analysis showed neurodevelopmental disorders as one of the largest and most statistically significant categories (Figure 3D). However, the other disease groups such as facial dysmorphisms, motor delay, cognitive problems or seizures (abnormal encephalogram) have also been associated multiple neurodevelopmental disorders and syndromic ASD ^47^. To further understand the molecular signatures associated with DTU related neurodevelopmental disorders we examined the genes in this category (Figure 3E). We found an enrichment of genes that have high risk variants associated with ASD (i.e. *CHD8*, *FOXP1*, *SHANK3*) ^48^ and broader neuronal development (i.e. *ROBO1*, *NDE1*, *RACK1*) ^49–52^. Taken together, our work suggests that regulation of DTU might be essential for neurodevelopment.

**Figure 3:**
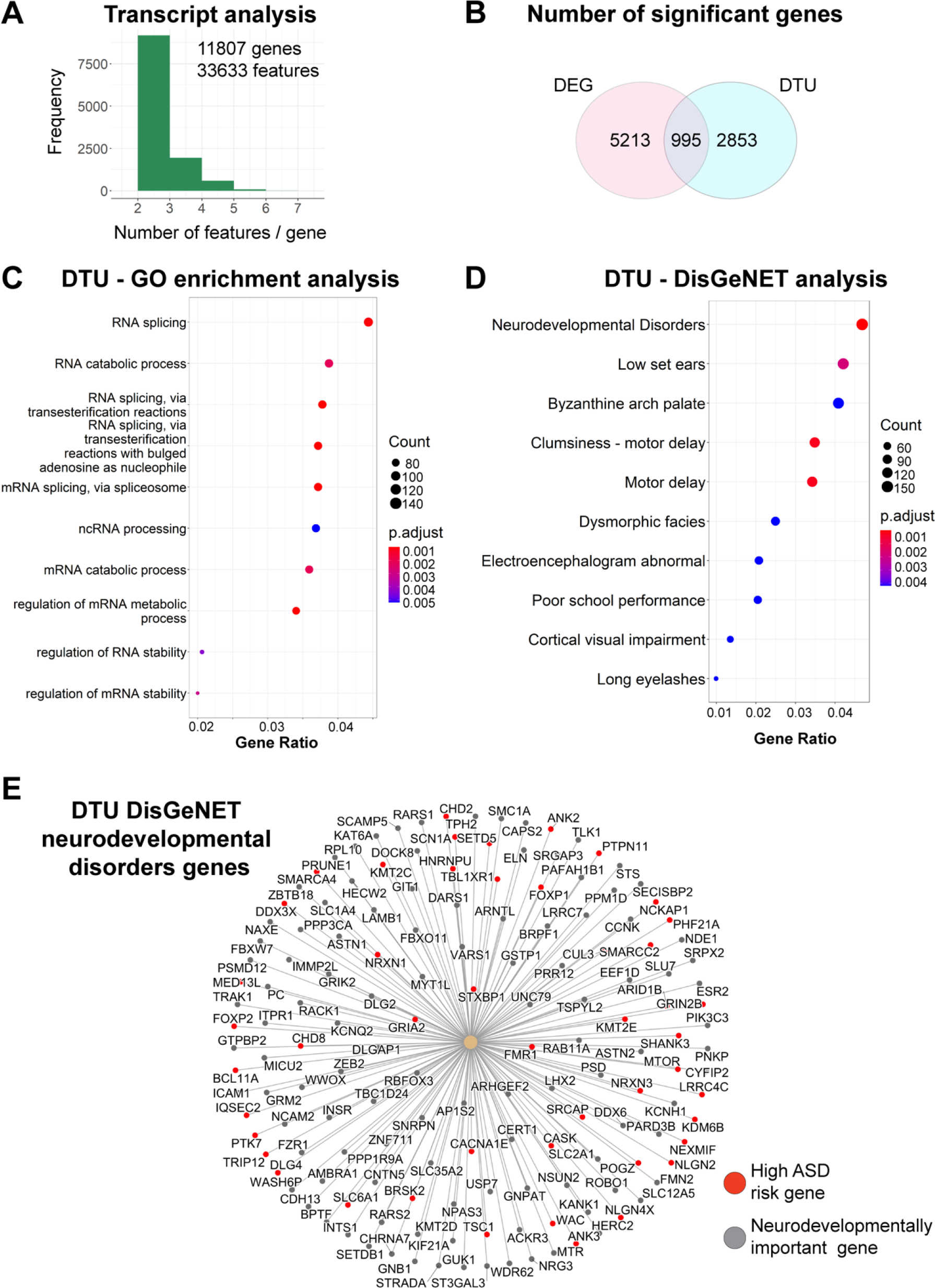
Differential transcript usage analysis reveals differential transcript usage (DTU) of genes associated with RNA splicing mechanisms and overrepresentation of genes associated with neurodevelopmental disorders. **(A)** Bar plot showing total number of genes and transcript isoforms analyzed in DRIMSeq DTU analysis in VPA and control samples. **(B)** Venn diagram showing overlap between significant differentially expressed genes (DEGs) (LFC > |1.5| and adjusted *P* < 0.05) and significant differential transcript usage (DTU) events (*P* < 0.05). **(C)** Gene ontology (GO) analysis of significant DTU events (*q* value cutoff = 0.05). Top 10 enriched pathways are shown. **(D)** Characterization of disease pathways by DisGeNET analysis of significant DTU events. Top 10 enriched pathways are shown. **(E)** Network plot of genes showing significant DTU events identified in the neurodevelopmental disorder DisGeNET pathway. High confidence ASD risk genes (category 1 in SFARI gene database) are labeled in red. Neurodevelopmentally important genes are labeled in gray.

### VPA differentially regulates gene expression and transcript usage of ASD-risk genes

The identification of ASD-risk genes among genes with DTU prompted us to define the extent to which ASD-risk genes overlapped with DEGs and DTU genes in VPA treated neurons. We found that close to 25% of either DEGs or DTU genes were ASD-risk factors (Figure 4A). However, only one fourth of the ASD-risk genes among the DEGs and DTU genes overlapped (Figure 4B). We performed GO analysis to determine whether the DEG- or DTU-specific genes in the ASD-risk gene group were enriched for different pathways (Supplementary Table S9). Top pathways revealed from GO biological process analysis of both DEG- and DTU-specific genes included: modulation of chemical synaptic transmission, regulation of post-synaptic signaling, and synapse organization (Supplementary Figure S4). GO cellular component analysis of both DEG- and DTU-specific genes revealed enrichment in synaptic membrane and pre-synapse pathways (Supplementary Figure S4). Interestingly, GO molecular function analysis revealed that DEG- and DTU-specific genes were enriched in different molecular function pathways. DEG-specific genes were enriched for ion channel activity and transmembrane transporter activity (Figure 4C). However, DTU-specific genes were enriched for histone binding, RNA II polymerase binding, and transcription machinery binding (Figure 4D). We further analyzed how DTU was altering specific genes, by focusing on two representative ASD-risk genes *SETD5* and *SMARCA4* (Figure 4E). Analysis of *SETD5* transcripts showed that VPA treated neurons had decreased levels of a full-length protein coding isoform, and increased levels of two non-coding isoforms. In contrast, analysis of *SMARCA4* transcripts in VPA treated neurons showed increased levels of the largest protein coding isoform, but decreased levels of a shorter protein coding isoform and a short, processed transcript. In summary, our analysis of ASD-risk genes dysregulated in VPA-treated neurons suggests that DEGs are enriched with genes associated with the control of synaptic organization and function, while genes with DTU are enriched with genes associated with control of transcription and chromatin architecture.

**Figure 4:**
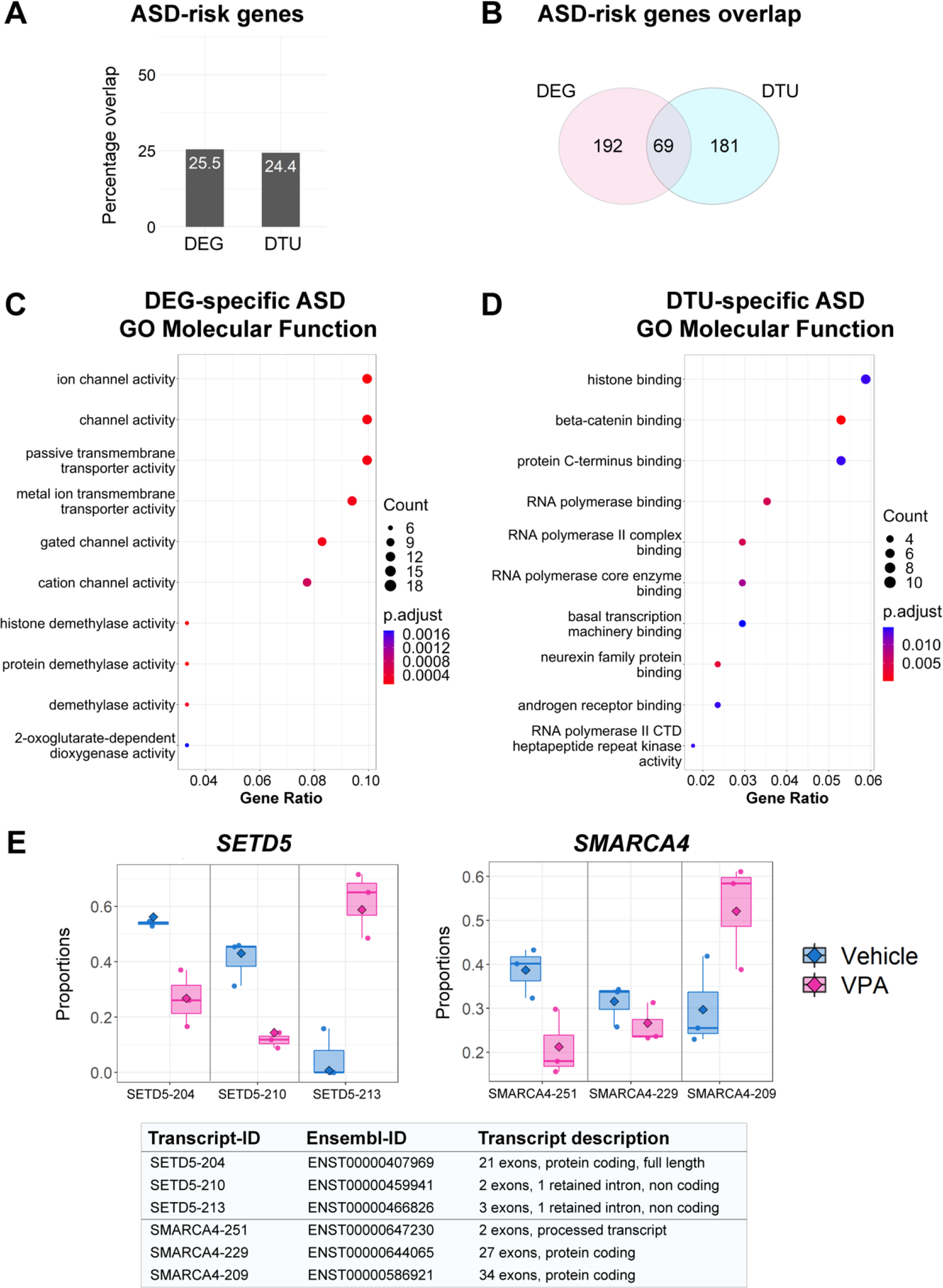
ASD-risk genes are differentially affected by gene expression and transcript usage in VPA-treated neurons. **(A)** Bar plot showing overlap between significant DEGs (LFC > |1.5| and adjusted *P* < 0.05) or DTU events (*P* < 0.05) with ASD-risk genes. **(B)** Venn diagram showing overlap of ASD-risk genes between significant DEGs and significant DTU events. **(C)** Gene ontology (GO) molecular function (MF) analysis of DEG-specific ASD-risk genes. **(D)** GO MF analysis of DTU-specific ASD-risk genes. **(E)** Analysis of two representative genes with DTUs. Box plots are shown for different transcripts in chromatin regulators *SETD5* (left panel) and *SMARCA4* (right panel). Transcripts found in vehicle treated neurons are shown in blue, and transcripts found in VPA treated neurons are shown in magenta. Description of different transcripts and ensemble IDs are shown in bottom panel.

## DISCUSSION

A recent study on non-mature human forebrain organoids treated with VPA for 3 days showed primarily alterations in synaptic transmission and ASD-risk genes which correlates with our own findings ^53^. However, in our study, we find that these same genes, in addition to other genes encoding chromatin, transcription, and splicing regulators, display differential transcript usage, which may amplify VPA’s effect on gene expression. Previously, a transcriptomic study in rats showed that there were significant genome-wide changes in differential splicing in response to VPA ^54^. Similarly, gene-specific changes in differential exon usage have been reported for *BDNF* in mice treated with VPA in utero ^55^. We also find that VPA elicits differential gene expression of genes associated with RNA splicing, as well as genome-wide changes in RNA splicing, which is a unique signature that had not been previously reported in human neurons in response to VPA. Taken together, these data suggest that the effect of VPA in neuronal development might be mediated by changes not only on gene expression, but potentially in the control of alternative splicing mechanisms.

The post-translational acetylation of histones is a well-known mechanism associated with activation of transcription ^56^. Treatment with VPA, a potent HDAC inhibitor, results in a hyperacetylated chromatin environment, which mitigates HDAC-dependent transcriptional repression ^13,14^. In addition, the regulation of histone acetylation provides a broad mechanism to regulate alternative splicing and these mechanisms has been described in neurons ^57^ and astrocytes ^58^. Different HDAC enzymes have been shown to interact with different splicing factors ^22^, and the activity of HDACs is proposed to influence splice site selection ^21^. Histone hyperacetylation has also been shown to disrupt physical interaction between splicing factors and chromatin. For instance, SF3B1, a component of the U2 snRNP, was shown to physically associate with chromatin and facilitate splice-site recognition ^59^. Interestingly, addition of VPA was found to disrupt the association between SF3B1 and chromatin and led to decreased exon inclusion levels on *SF3B1*-enriched exons in HeLa cells ^59^. Therefore, we posit that by inducing a hyperacetylated chromatin environment through its HDAC inhibitory function, VPA is altering not just gene expression, but also RNA splicing (Figure 5). Support for this hypothesis is provided by the identification of a general downregulation in the types of splicing events as well as a unique molecular signature of genes with DTU in VPA-treated neurons. We observed reduced numbers of alternative exon inclusion events in the VPA-treated neurons. This finding coincides with the downregulation of *RBFOX2*, and the presence of DTU in *RBFOX3* in response to VPA. RBFOX genes encode splicing regulatory factors that have been associated with either promoting or repressing exon inclusion during splicing ^60^. Therefore, the distinct dysregulation of both *RBFOX2* and *RBFOX3*, among other genes encoding splicing factors, could contribute in part to the overall alterations in RNA splicing associated with VPA exposure.

**Figure 5:**
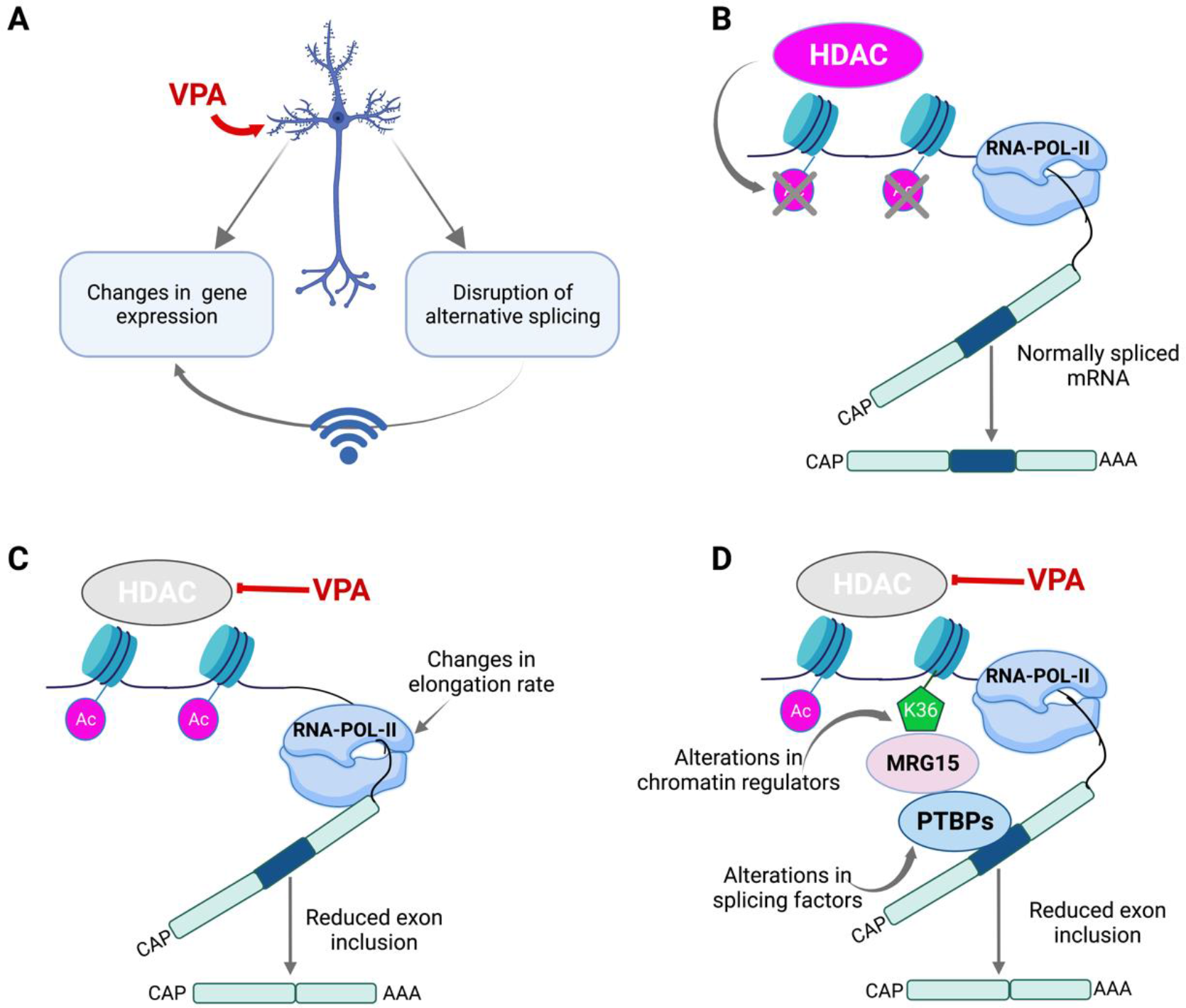
Potential models of VPA-induced changes in alternative splicing and differential transcript usage in human neurons. **(A)** Summary on the effect of VPA in human neurons. VPA elicits changes in gene expression and mRNA splicing. The effect of VPA in mRNA splicing could in turn further exacerbate VPA’s effect in the changes of gene expression. **(B)** Normal HDAC function, and the coupling of transcription by RNA polymerase II (RNA pol-II) and pre-mRNA splicing is shown. **(C)** Diagram shows changes in RNA elongation rate influenced by increased histone acetylation in VPA treated neurons leading to defects in pre-mRNA splicing. **(D)** Diagram shows that alterations VPA exposure could lead to alterations in the splicing machinery or chromatin regulators that constitute interacting hubs for splicing factors, leading to alternative splicing defects.

The influence of the chromatin environment in the co-transcriptional regulation of RNA splicing is achieved by numerous mechanisms that implicate chromatin remodeling and post-translational modifications of histones ^61^. Our work shows that expression of genes related to histone and chromatin regulators were significantly affected, and this could additionally impact splicing outcomes in response to VPA. One well-studied example of a histone modification that regulates RNA splicing is methylation of H3K36, a highly conserved histone modification. In yeast, it was found that H3K36 methylation was required for proper co-transcriptional spliceosome assembly by recruiting a chromodomain protein that stabilized interaction between splicing factors and chromatin ^62^. In humans, H3K36me3 was implicated in regulating alternative splicing via recruitment of polypyrimidine tract-binding protein (PTB) via chromodomain protein MRG15 ^61^. We find evidence that VPA modulates genes encoding histone methyltransferases that methylate H3K36, such as NSD1 ^63^ and SETD5 ^64^. NSD1 has been associated with syndromic ASD ^65^, and modulates the mono- and di-methylation of H3K36 (H3K36me1/2) ^63^. Human neurons treated with VPA showed significantly decreased expression of *NSD1* in this study. H3K36me2 serves as a substrate for the subsequent tri-methylation of H3K36. Therefore, loss of NSD1 could lead to a widespread reduction of H3K36me3 at a genome-wide scale, which in turn could disrupt alternative splicing. Similarly, we find that SETD5, which catalyzes the trimethylation of H3K36 ^64^, shows DTU with decreased levels of the protein coding isoform and increased levels of two non-coding isoforms. Mice haploinsufficient for SETD5 have decrease neuronal progenitor proliferation, synaptic defects, decreased sociability and impairments on cognitive tasks, which correlates with loci-specific reduction of H3K36me3 ^64^, and increased chromatin hyperacetylation ^66^. Taken together, these data suggest that in human neurons exposure to VPA could reduce the expression of *NSD1* and the coding *SETD5* isoform and lead to decreased H3K36me3 levels, which in turn can affect RNA splicing. Therefore, we posit that dysregulation of H3K36 methylation, as well as potentially dysregulation of other histone modifications ^61,67^, could alter the chromatin environment further exacerbating the alternative splicing defects associated with the hyperacetylation of chromatin caused by VPA exposure.

The relevance of our findings on the dysregulation of alternative splicing to the mechanisms that underlie ASD pathogenesis is suggested by its correlation with previous transcriptomic studies of ASD-postmortem brain tissue. A study of over 1600 post-mortem cerebral cortex samples from ASD, schizophrenia (SCZ), and bipolar disorder (BP) individuals showed that across the three disorders changes in isoform-level had the largest effect size associated with disease ^68^. Moreover, like our own findings in which VPA induces enrichment of splicing regulatory proteins among genes with DTU, analysis of ASD, SCZ, and BP cerebral cortices also showed disease association in the splicing of RNA binding proteins and splicing factors ^68^. Further a study of 48 ASD-postmortem cerebral cortex tissue, showed downregulation of genes involved in the control of alternative splicing ^69^. We present three possible mechanisms by which chromatin dysregulation by VPA affects RNA splicing in human neurons that could potentially contribute to ASD pathogenesis (Figure 5). Further studies are necessary to precisely dissect the role of VPA in the regulation of RNA splicing. In summary, our work highlights the importance of the chromatin environment in the regulation of alternative splicing in the pathogenesis of ASD associated with environmental risk factors and potentially genetic risk factors (chromatin regulators).

## ACKNOWLEDGEMENTS

We thank Dr. Alper Uzun, and members of the Lizarraga laboratory for critical feedback on the manuscript. RNA sequencing was conducted using the molecular core at Brown University. qPCR experiments were carried out using the molecular core at the center of childhood neurotherapeutics at UofSC. SBL was supported by an NIH/NIMH R01 MH127081-01A1, NIH/NIGMS COBRE 4P20GM103641-06, SC EPSCOR stimulus award 18-SR04, NIH/NIGMS SC INBRE pilot 5P20GM103499-17, and ASPIRE awards Track I and Track II from the VPR office at UofSC. EDGU was supported by Brown University Legorreta Cancer Center and Rhode Island Foundation. CSL was supported by T32 postdoctoral grant #.

## REFERENCES

1 Gandal, M. J., Leppa, V., Won, H., Parikshak, N. N. & Geschwind, D. H. The road to precision psychiatry: translating genetics into disease mechanisms. Nat Neurosci 19, 1397–1407, doi:10.1038/nn.4409 (2016).

2 Modabbernia, A., Velthorst, E. & Reichenberg, A. Environmental risk factors for autism: an evidence-based review of systematic reviews and meta-analyses. Mol Autism 8, 13, doi:10.1186/s13229-017-0121-4 (2017).

3 Chaste, P. & Leboyer, M. Autism risk factors: genes, environment, and gene-environment interactions. Dialogues Clin Neurosci 14, 281–292 (2012).

4 Rasalam, A. D. et al. Characteristics of fetal anticonvulsant syndrome associated autistic disorder. Dev Med Child Neurol 47, 551–555, doi:10.1017/s0012162205001076 (2005).

5 Christensen, J. et al. Prenatal valproate exposure and risk of autism spectrum disorders and childhood autism. JAMA 309, 1696–1703, doi:10.1001/jama.2013.2270 (2013).

6 Bromley, R. L. et al. The prevalence of neurodevelopmental disorders in children prenatally exposed to antiepileptic drugs. J Neurol Neurosurg Psychiatry 84, 637–643, doi:10.1136/jnnp-2012-304270 (2013).

7 Bromley, R. L. et al. Autism spectrum disorders following in utero exposure to antiepileptic drugs. Neurology 71, 1923–1924, doi:10.1212/01.wnl.0000339399.64213.1a (2008).

8 Moore, S. J. et al. A clinical study of 57 children with fetal anticonvulsant syndromes. J Med Genet 37, 489–497, doi:10.1136/jmg.37.7.489 (2000).

9 Mutlu-Albayrak, H., Bulut, C. & Caksen, H. Fetal Valproate Syndrome. Pediatr Neonatol 58, 158–164, doi:10.1016/j.pedneo.2016.01.009 (2017).

10 Williams, G. et al. Fetal valproate syndrome and autism: additional evidence of an association. Dev Med Child Neurol 43, 202–206 (2001).

11 Struhl, K. Histone acetylation and transcriptional regulatory mechanisms. Genes Dev 12, 599–606, doi:10.1101/gad.12.5.599 (1998).

12 Kuo, M. H. & Allis, C. D. Roles of histone acetyltransferases and deacetylases in gene regulation. Bioessays 20, 615–626, doi:10.1002/(SICI)1521-1878(199808)20:8<615∷AID-BIES4>3.0.CO;2-H (1998).

13 Kramer, O. H. et al. The histone deacetylase inhibitor valproic acid selectively induces proteasomal degradation of HDAC2. EMBO J 22, 3411–3420, doi:10.1093/emboj/cdg315 (2003).

14 Gottlicher, M. et al. Valproic acid defines a novel class of HDAC inhibitors inducing differentiation of transformed cells. EMBO J 20, 6969–6978, doi:10.1093/emboj/20.24.6969 (2001).

15 Kataoka, S. et al. Autism-like behaviours with transient histone hyperacetylation in mice treated prenatally with valproic acid. Int J Neuropsychopharmacol 16, 91–103, doi:10.1017/S1461145711001714 (2013).

16 Drogaris, P. et al. Histone deacetylase inhibitors globally enhance h3/h4 tail acetylation without affecting h3 lysine 56 acetylation. Sci Rep 2, 220, doi:10.1038/srep00220 (2012).

17 Slaughter, M. J. et al. HDAC inhibition results in widespread alteration of the histone acetylation landscape and BRD4 targeting to gene bodies. Cell Rep 34, 108638, doi:10.1016/j.celrep.2020.108638 (2021).

18 Rahhal, R. & Seto, E. Emerging roles of histone modifications and HDACs in RNA splicing. Nucleic Acids Res 47, 4911–4926, doi:10.1093/nar/gkz292 (2019).

19 Gunderson, F. Q. & Johnson, T. L. Acetylation by the transcriptional coactivator Gcn5 plays a novel role in co-transcriptional spliceosome assembly. PLoS Genet 5, e1000682, doi:10.1371/journal.pgen.1000682 (2009).

20 Will, C. L. & Luhrmann, R. Spliceosome structure and function. Cold Spring Harb Perspect Biol 3, doi:10.1101/cshperspect.a003707 (2011).

21 Hnilicova, J. et al. Histone deacetylase activity modulates alternative splicing. PLoS One 6, e16727, doi:10.1371/journal.pone.0016727 (2011).

22 Khan, D. H. et al. RNA-dependent dynamic histone acetylation regulates MCL1 alternative splicing. Nucleic Acids Res 42, 1656–1670, doi:10.1093/nar/gkt1134 (2014).

23 Bromley, R. et al. Treatment for epilepsy in pregnancy: neurodevelopmental outcomes in the child. Cochrane Database Syst Rev, CD010236, doi:10.1002/14651858.CD010236.pub2 (2014).

24 Shallcross, R. et al. Child development following in utero exposure: levetiracetam vs sodium valproate. Neurology 76, 383–389, doi:10.1212/WNL.0b013e3182088297 (2011).

25 Reid, C. B. & Walsh, C. A. Early development of the cerebral cortex. Prog Brain Res 108, 17-30 (1996).

26 Manzini, M. C. & Walsh, C. A. What disorders of cortical development tell us about the cortex: one plus one does not always make two. Curr Opin Genet Dev 21, 333–339, doi:10.1016/j.gde.2011.01.006 (2011).

27 Mariani, J. et al. Modeling human cortical development in vitro using induced pluripotent stem cells. Proc Natl Acad Sci U S A 109, 12770–12775, doi:10.1073/pnas.1202944109 (2012).

28 Shi, Y., Kirwan, P., Smith, J., Robinson, H. P. & Livesey, F. J. Human cerebral cortex development from pluripotent stem cells to functional excitatory synapses. Nat Neurosci 15, 477–486, S471, doi:10.1038/nn.3041 (2012).

29 Kirwan, P. et al. Development and function of human cerebral cortex neural networks from pluripotent stem cells in vitro. Development 142, 3178–3187, doi:10.1242/dev.123851 (2015).

30 Shi, Y., Kirwan, P. & Livesey, F. J. Directed differentiation of human pluripotent stem cells to cerebral cortex neurons and neural networks. Nat Protoc 7, 1836–1846, doi:10.1038/nprot.2012.116 (2012).

31 van de Leemput, J. et al. CORTECON: a temporal transcriptome analysis of in vitro human cerebral cortex development from human embryonic stem cells. Neuron 83, 51–68, doi:10.1016/j.neuron.2014.05.013 (2014).

32 Abrahams, B. S. et al. SFARI Gene 2.0: a community-driven knowledgebase for the autism spectrum disorders (ASDs). Mol Autism 4, 36, doi:10.1186/2040-2392-4-36 (2013).

33 Parikshak, N. N. et al. Genome-wide changes in lncRNA, splicing, and regional gene expression patterns in autism. Nature 540, 423–427, doi:10.1038/nature20612 (2016).

34 Voineagu, I. et al. Transcriptomic analysis of autistic brain reveals convergent molecular pathology. Nature 474, 380–384, doi:10.1038/nature10110 (2011).

35 Lizarraga, S. B. et al. Human neurons from Christianson syndrome iPSCs reveal mutation-specific responses to rescue strategies. Sci Transl Med 13, doi:10.1126/scitranslmed.aaw0682 (2021).

36 Cheon, S. et al. Counteracting epigenetic mechanisms regulate the structural development of neuronal circuitry in human neurons. Mol Psychiatry, doi:10.1038/s41380-022-01474-1 (2022).

37 Andrews, S. FastQC: A Quality Control Tool for High Throughput Sequence Data, <https://www.bioinformatics.babraham.ac.uk/projects/fastqc/> (2010).

38 Patro, R., Duggal, G., Love, M. I., Irizarry, R. A. & Kingsford, C. Salmon provides fast and bias-aware quantification of transcript expression. Nat Methods 14, 417–419, doi:10.1038/nmeth.4197 (2017).

39 Soneson, C., Love, M. I. & Robinson, M. D. Differential analyses for RNA-seq: transcript-level estimates improve gene-level inferences. F1000Res 4, 1521, doi:10.12688/f1000research.7563.2 (2015).

40 Love, M. I., Huber, W. & Anders, S. Moderated estimation of fold change and dispersion for RNA-seq data with DESeq2. Genome Biol 15, 550, doi:10.1186/s13059-014-0550-8 (2014).

41 Nowicka, M. & Robinson, M. D. DRIMSeq: a Dirichlet-multinomial framework for multivariate count outcomes in genomics. F1000Res 5, 1356, doi:10.12688/f1000research.8900.2 (2016).

42 Shen, S. et al. rMATS: robust and flexible detection of differential alternative splicing from replicate RNA-Seq data. Proc Natl Acad Sci U S A 111, E5593–5601, doi:10.1073/pnas.1419161111 (2014).

43 Willsey, A. J. et al. Coexpression networks implicate human midfetal deep cortical projection neurons in the pathogenesis of autism. Cell 155, 997–1007, doi:10.1016/j.cell.2013.10.020 (2013).

44 Lee, Y. & Rio, D. C. Mechanisms and Regulation of Alternative Pre-mRNA Splicing. Annu Rev Biochem 84, 291–323, doi:10.1146/annurev-biochem-060614-034316 (2015).

45 Tazi, J., Bakkour, N. & Stamm, S. Alternative splicing and disease. Biochim Biophys Acta 1792, 14–26, doi:10.1016/j.bbadis.2008.09.017 (2009).

46 Pinero, J. et al. DisGeNET: a comprehensive platform integrating information on human disease-associated genes and variants. Nucleic Acids Res 45, D833–D839, doi:10.1093/nar/gkw943 (2017).

47 Hanly, C., Shah, H., Au, P. Y. B. & Murias, K. Description of neurodevelopmental phenotypes associated with 10 genetic neurodevelopmental disorders: A scoping review. Clin Genet 99, 335–346, doi:10.1111/cge.13882 (2021).

48 Satterstrom, F. K. et al. Large-Scale Exome Sequencing Study Implicates Both Developmental and Functional Changes in the Neurobiology of Autism. Cell 180, 568–584 e523, doi:10.1016/j.cell.2019.12.036 (2020).

49 Gonda, Y. et al. Robo1 regulates the migration and laminar distribution of upper-layer pyramidal neurons of the cerebral cortex. Cereb Cortex 23, 1495–1508, doi:10.1093/cercor/bhs141 (2013).

50 Andrews, W. et al. Robo1 regulates the development of major axon tracts and interneuron migration in the forebrain. Development 133, 2243–2252, doi:10.1242/dev.02379 (2006).

51 Alkuraya, F. S. et al. Human mutations in NDE1 cause extreme microcephaly with lissencephaly [corrected]. Am J Hum Genet 88, 536–547, doi:10.1016/j.ajhg.2011.04.003 (2011).

52 Zhu, Q. et al. Rack1 is essential for corticogenesis by preventing p21-dependent senescence in neural stem cells. Cell Rep 36, 109639, doi:10.1016/j.celrep.2021.109639 (2021).

53 Meng, Q. et al. Human forebrain organoids reveal connections between valproic acid exposure and autism risk. Transl Psychiatry 12, 130, doi:10.1038/s41398-022-01898-x (2022).

54 Zhang, R. et al. Transcriptional and splicing dysregulation in the prefrontal cortex in valproic acid rat model of autism. Reprod Toxicol 77, 53–61, doi:10.1016/j.reprotox.2018.01.008 (2018).

55 Konopko, M. A., Densmore, A. L. & Krueger, B. K. Sexually Dimorphic Epigenetic Regulation of Brain-Derived Neurotrophic Factor in Fetal Brain in the Valproic Acid Model of Autism Spectrum Disorder. Dev Neurosci 39, 507–518, doi:10.1159/000481134 (2017).

56 Imhof, A. & Wolffe, A. P. Transcription: gene control by targeted histone acetylation. Curr Biol 8, R422–424, doi:10.1016/s0960-9822(98)70268-4 (1998).

57 Zhou, H. L. et al. Hu proteins regulate alternative splicing by inducing localized histone hyperacetylation in an RNA-dependent manner. Proc Natl Acad Sci U S A 108, E627–635, doi:10.1073/pnas.1103344108 (2011).

58 Kanski, R. et al. Histone acetylation in astrocytes suppresses GFAP and stimulates a reorganization of the intermediate filament network. J Cell Sci 127, 4368–4380, doi:10.1242/jcs.145912 (2014).

59 Kfir, N. et al. SF3B1 association with chromatin determines splicing outcomes. Cell Rep 11, 618–629, doi:10.1016/j.celrep.2015.03.048 (2015).

60 Lovci, M. T. et al. Rbfox proteins regulate alternative mRNA splicing through evolutionarily conserved RNA bridges. Nat Struct Mol Biol 20, 1434–1442, doi:10.1038/nsmb.2699 (2013).

61 Luco, R. F. et al. Regulation of alternative splicing by histone modifications. Science 327, 996–1000, doi:10.1126/science.1184208 (2010).

62 Leung, C. S. et al. H3K36 Methylation and the Chromodomain Protein Eaf3 Are Required for Proper Cotranscriptional Spliceosome Assembly. Cell Rep 27, 3760–3769 e3764, doi:10.1016/j.celrep.2019.05.100 (2019).

63 Lucio-Eterovic, A. K. et al. Role for the nuclear receptor-binding SET domain protein 1 (NSD1) methyltransferase in coordinating lysine 36 methylation at histone 3 with RNA polymerase II function. Proc Natl Acad Sci U S A 107, 16952–16957, doi:10.1073/pnas.1002653107 (2010).

64 Sessa, A. et al. SETD5 Regulates Chromatin Methylation State and Preserves Global Transcriptional Fidelity during Brain Development and Neuronal Wiring. Neuron 104, 271–289 e213, doi:10.1016/j.neuron.2019.07.013 (2019).

65 Muhsin, E., Basak, G., Banu, D., Alper, G. & Mustafa, S. Neurodevelopment and Genetic Evaluation of Sotos Syndrome Cases with a Novel Mutation: a Single-Center Experience. J Mol Neurosci 72, 149–157, doi:10.1007/s12031-021-01897-5 (2022).

66 Deliu, E. et al. Haploinsufficiency of the intellectual disability gene SETD5 disturbs developmental gene expression and cognition. Nat Neurosci 21, 1717–1727, doi:10.1038/s41593-018-0266-2 (2018).

67 Schor, I. E., Fiszbein, A., Petrillo, E. & Kornblihtt, A. R. Intragenic epigenetic changes modulate NCAM alternative splicing in neuronal differentiation. EMBO J 32, 2264–2274, doi:10.1038/emboj.2013.167 (2013).

68 Gandal, M. J. et al. Transcriptome-wide isoform-level dysregulation in ASD, schizophrenia, and bipolar disorder. Science 362, doi:10.1126/science.aat8127 (2018).

69 Parikshak, N. N. et al. Genome-wide changes in lncRNA, splicing, and regional gene expression patterns in autism. Nature, doi:10.1038/nature20612 (2016).

