## Supplementary Material for "Dysregulation of The Chromatin Environment Leads to Differential Alternative Splicing as A Mechanism Of Disease In a Human Model of Autism Spectrum Disorders"

Calvin S. Leung ^1, 2^, Shoshanna Rosenzweig ^3, 4^, Brian Yoon ^1, 2,^ Nicholas A. Marinelli ^1, 2^, Ethan W. Hollingsworth ^5, 6^, Abbie M. Maguire ^7, 8^, Mara M. Cowen ^1, 2^, Michael Schmidt ^7, 8^, Jaime Imitola ^5, 6^, Ece D. Gamsiz Uzun ^3. 4, &^ and Sofia B. Lizarraga^1, 2, &^

^1^ Department of Biological Sciences, University of South Carolina, 715 Sumter Street, Columbia, SC 29208

^2^ Center for Childhood Neurotherapeutics, University of South Carolina, 715 Sumter Street, Columbia, SC 29208

^3^ Department of Pathology and Laboratory Medicine, Alpert Medical School of Brown University, Providence, RI 02912

^4^ Center for Computational Molecular Biology, Brown University, Providence, RI 02906

^5^ UCONN Health Comprehensive Multiple Sclerosis Center, Department of Neurology, University of Connecticut School of Medicine, Farmington, CT 06030

^6^ Division of Multiple Sclerosis and Translational Neuroimmunology, Department of Neurology, University of Connecticut School of Medicine, Farmington, CT 06030

^7^ Department of Molecular Biology, Cell Biology and Biochemistry; Institute for Brain Science; and the Hassenfeld Child Health Innovation Institute, Brown University, Lab for Molecular Medicine, 70 Ship Street, Providence, RI 02912

^8^ Developmental Disorders Genetics Research Program, Emma Pendleton Bradley Hospital and Department of Psychiatry and Human Behavior, Alpert Medical School of Brown University, 1011 Veteran Memorial Pkwy. East Providence, RI 02912

^&^ Co-corresponding authors

**Running Title:** Co-regulation of chromatin environment and alternative splicing in autism pathogenesis

**Keywords:** autism, valproic acid, stem cells, chromatin, alternative splicing, isoforms, hyperacetylation

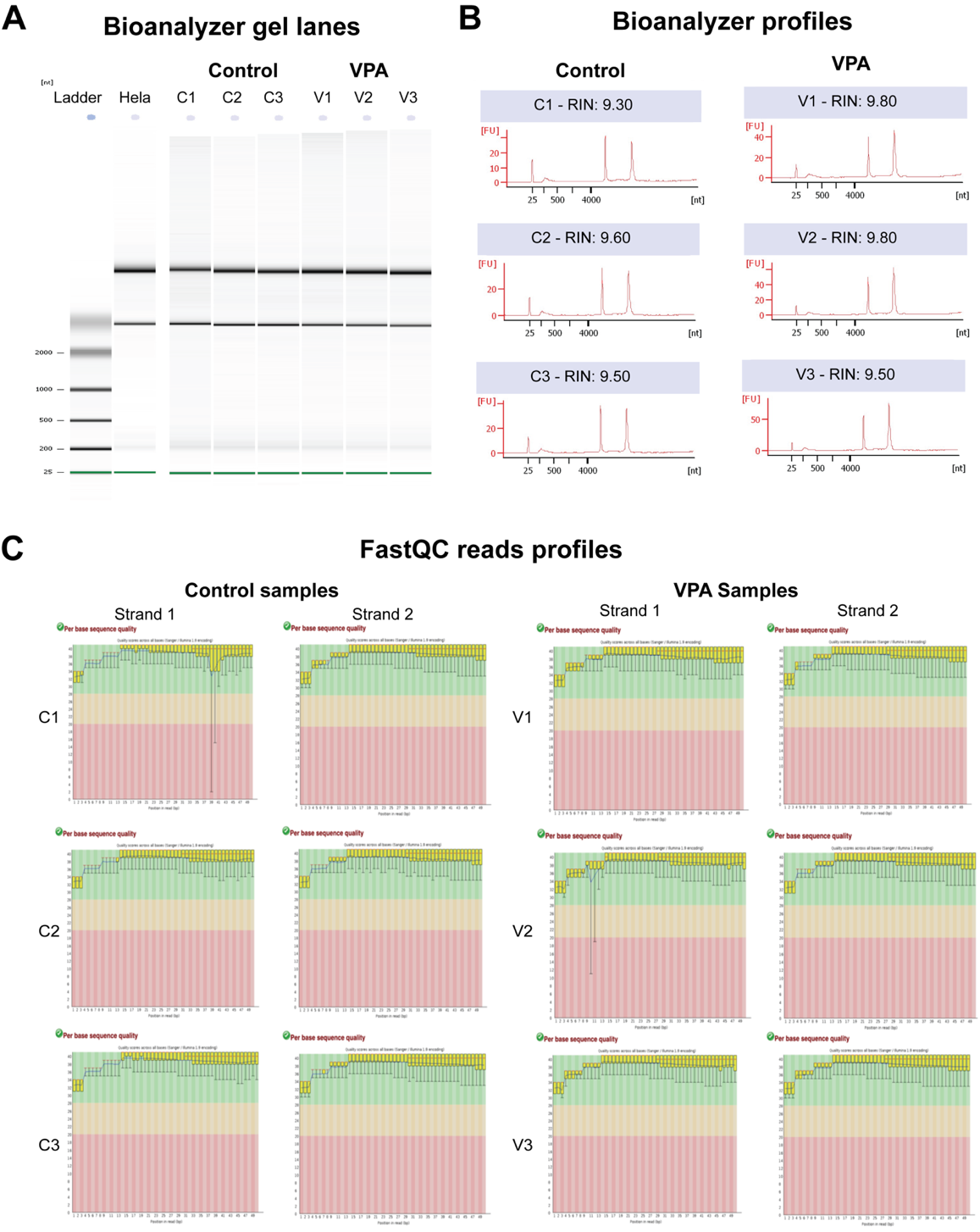
**SUPPLEMENTARY FIGURES AND FIGURE LEGENDS**

**Supplementary Figure S1: Quality control for total RNA and RNA-seq reads. (A-B)** Analysis of total RNA quality by bioanalyzer. RNA bands (**A**) as well as the bioanalyzer profiles with RNA integrity number (RIN) (**B**) are shown for all control (vehicle) and VPA treated samples. (**C**) Fast-QC profiles show the quality of the RNA-reads in each RNA library for both strands in each control or VPA treated sample.

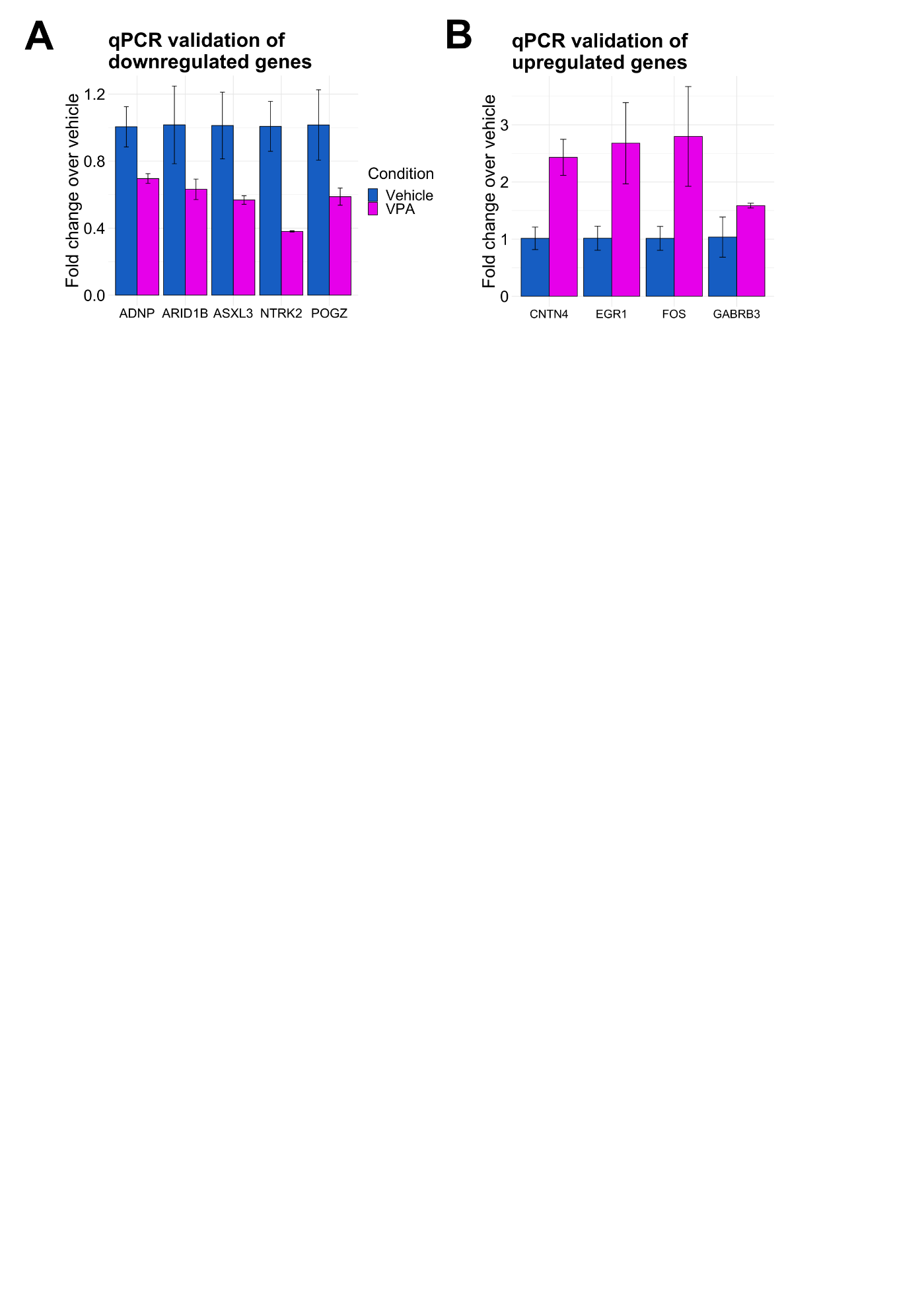
**Supplementary Figure S2: Gene expression validation of DESeq2 results. (A)** Gene expression analysis by qPCR of representative significantly downregulated DEGs in vehicle (blue) and VPA (magenta) treated neurons. **(B)** Gene expression analysis by qPCR of representative significantly upregulated DEGs, in vehicle (blue) and VPA treated neurons. Error bars represent standard deviation.

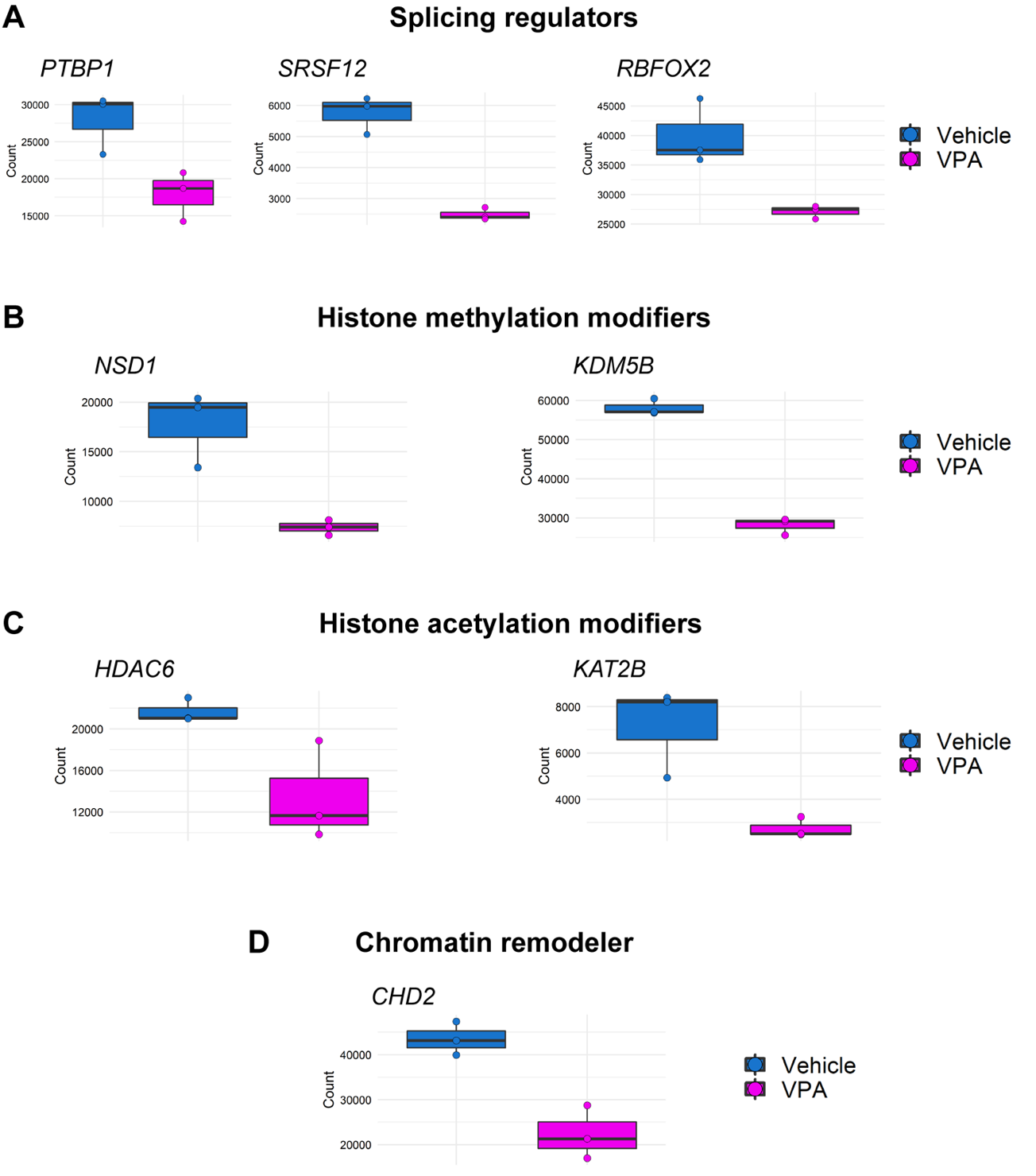

**Supplementary Figure S3: Normalized DESeq2 gene counts of genes identified in GSEA analysis. (A)** Normalized gene counts of representative alternative splicing factors identified in the regulation mRNA processing pathway in vehicle (blue) and VPA (magenta) samples. **(B)** Normalized gene counts of representative genes identified in the covalent chromatin modification, histone modification, and DNA conformation change pathways associated with the control of histone methylation in vehicle (blue) and VPA (magenta) samples. **(C)** Normalized gene counts of representative genes identified in the covalent chromatin modification, histone modification, and DNA conformation change pathways associated with the control of histone acetylation in vehicle (blue) and VPA (magenta) samples. **(D)** Normalized gene counts of representative genes identified in the covalent chromatin modification, histone modification, and DNA conformation change pathways associated with the control of chromatin remodeling in vehicle (blue) and VPA (magenta) samples. Error bars represent standard deviation.

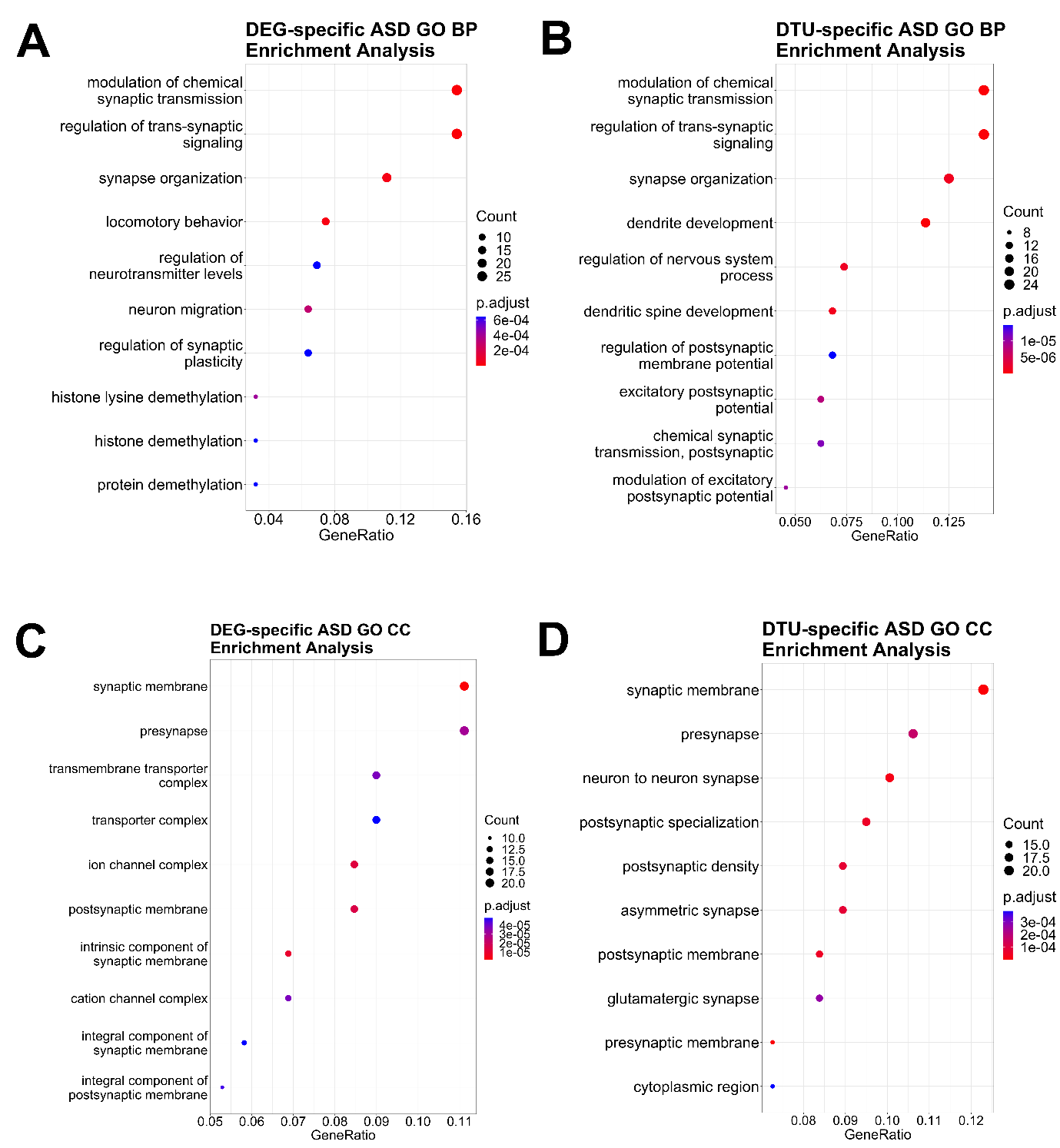

**Supplementary Figure S4: Gene ontology (GO) biological process (BP) and cellular component (CC) analysis of DEG- and DTU-specific ASD-risk genes. A.** GO BP analysis of DEG-specific ASD-risk genes. Top 10 enrichment pathways are shown. *P* value cutoff = 0.05. **B.** GO BP analysis of DTU-specific ASD-risk genes. Top 10 enrichment pathways are shown. *P* value cutoff = 0.05. **C.** GO CC analysis of DEG-specific ASD-risk genes. Top 10 enrichment pathways are shown. *P* value cutoff = 0.05. **D.** GO CC analysis of DTU-specific ASD-risk genes. Top 10 enrichment pathways are shown. *P* value cutoff = 0.05.

**LEGENDS FOR SUPPLEMENTARY TABLES**

All supplementary tables are in an excel format in dataset1.

**Supplementary Table S1: Information on Sequencing Reads.** Table shows the total number of fragments analyzed and the percentage of mapping for each sample.

**Supplementary Table S2: Differential gene expression analysis.** Table shows downregulated and upregulated DEGs detected with DESeq2 software package (version 1.32.0) ^1^ using an adjusted *P* < 0.05 and a fold change ≥ |1.5|. Fold change is shown in logarithmic scale, the standard error estimate for the log2 gold change is shown as lfcSE, the regular and adjusted P values are also shown.

**Supplementary Table S3: Primers for qPCR analysis.** Table shows the primers used for gene expression analysis. Primers were pre-designed to cover all isoforms for each gene. Forward and Reverse primer sequences are shown.

**Supplementary Table S4: Gene Set Enrichment Analysis.** Table shows all the categories identified by Gene Set Enrichment Analysis (GSEA) database. The total number of genes in each category are shown in the setSize column. Gene ontology identification numbers (ID) are shown for each category and the genes included in each category are shown under the core_enrichment column and are listed by their Ensembl IDs. The normalized enrichment score (NES) is calculated by GSEA and is corrected for multiple hypothesis testing (FDR). The upregulated categories have a positive enrichment score or NES and the downregulated categories have a negative enrichment score or NES.

**Supplementary Table S5: Analysis of alternative splicing events using rMATs.** Separate tables are shown for each type of alternative splicing event. The events analyzed include: Skipped exon (SE), retained intron (RI), alternative 5’ or 3′ splice site (A5SS or A3SS) and mutually exclusive exon (MXE). Genes are shown by gene symbol, with annotations for: chromosome, strand orientation, exon start (ES), exon end (EE) locations, false discovery rate (FDR), and inclusion level (IncLevel) is shown as well as the inclusion level difference (IncLevelDifference).

**Supplementary Table S6: Differential transcript usage analysis using DRIMseq.** Table shows all genes with DTU. The Gene column represents the FDR of whether that particular gene undergoes DTU. The Transcript column represents whether that particular transcript of the gene shows differential expression levels compared to other transcripts of that same gene.

**Supplementary Table S7: Gene ontology analysis of significant DTU events.** Table shows gene ontology analysis for genes with DTU events. The gene ontology (GO) categories analyzed were molecular function (MF), biological process (BP), and cellular compartment (CC) for all the different genes. GeneRatio represents the number of significant genes found in a GO term / total number of significant genes. BgRatio represents the number of genes in a GO term / total number of background genes. Genes for each category are shown by their Ensembl ID.

**Supplementary Table S8: Analysis of disease pathways across significant DTU events using DisGene databases.** Table shows analysis of disease pathways each pathway has a unique GO identifier. Genes in each category are shown by their gene symbol. Gene ratio and Bg ratio are shown along with normal and adjusted p values.

**Supplementary Table S9: Gene ontology analysis of ASD-risk genes with specific differential gene expression or differential transcript usage.** Tables show GO analysis for ASD-risk genes with either DEG (DEG-ASD) or DTU events (DTU-ASD). The gene ontology (GO) categories analyzed were molecular function (MF), biological process (BP), and cellular compartment (CC). GeneRatio represents the number of significant genes found in a GO term with respect to the total number of significant genes. BgRatio represents the number of genes in a GO term with respect to the total number of background genes. Genes for each category are shown by the Ensembl ID.

**REFERENCES**

1	Love, M. I., Huber, W. & Anders, S. Moderated estimation of fold change and dispersion for RNA-seq data with DESeq2. *Genome Biol* **15**, 550, doi:10.1186/s13059-014-0550-8 (2014).
